# Testing an alternative explanation for relatively greater base-sharing between Neanderthals and non-African humans

**DOI:** 10.1101/133306

**Authors:** William Amos

## Abstract

Most accept that non-African humans share ∼2% of their genome with Neanderthals (1) and that inter-breeding occurred between several archaic lineages (2-4). However, most evidence assumes that mutation rate is constant. It has been suggested that heterozygosity is mutagenic (5-8). If so, an alternative explanation of the data becomes possible. Instead of non-Africans sharing relatively more bases with Neanderthals due to interbreeding, Africans could appear unexpectedly divergent due to their mutation rate not having been lowered when diversity was lost during the out of Africa bottleneck. I therefore tested a series of predictions aimed at distinguishing mutation slowdown from inter-breeding. Predictions from mutation slowdown are generally better supported. For example, the signal used to infer inter-breeding remains even when Neanderthal sequences are excluded. I conclude that, while some inter-breeding probably did occur, an appreciable component of the signal seems better explained by mutation slowdown.

## Introduction

The draft Neanderthal genome revealed more shared nucleotide bases with modern non-African humans compared with Africans (1). This asymmetric base-sharing was used to argue that Neanderthals inter-bred with the ancestors of modern non-Africans, leaving a modern genetic legacy of ∼2%. Subsequent studies have reinforced this model, with the discovery of skeletons with recent hybrid origin (9), an estimated 5% legacy from Denisovans in Oceania (10-12) and a decrease in the inferred size of introgressed blocks over time (13, 14), consistent with recombination breaking down introgressed blocks.

Inter-breeding with Neanderthals was originally inferred from four-way DNA sequence alignments comprising two humans, a Neanderthal and a chimpanzee (the ‘ABBA-BABA’ test (15-17)) (1, 16). Focus centres on bases where the Neanderthal and chimpanzee differ, with each matching one of the two humans (states ‘ABBA’ and ‘BABA’). Under a null model, with no introgression and constant mutation rate, ABBA and BABA sites should be equally frequent. In practice, when the two humans comprise one African and one non-African, the counts of ABBA and BABA are significantly asymmetrical (1). Such asymmetry, usually expressed as Patterson’s D (18), is most obviously explained by introgression of Neanderthal DNA into the non-African.

Subsequent studies have developed the ABBA-BABA into derivative statistics such as D_enhanced_ (10) and f4-ratios (18). Despite these developments, the underlying principles remain largely unchanged. Ultimately, all methods attempt to quantify excess base-sharing between an archaic genome and one modern genome relative to a control, usually a second modern human. Clusters of shared bases have been interpreted as introgressed haplotypes [12, 19]. However, many aspects of the mutation process remain poorly understood, including the lability and strength of mutation hotspots (19), the tendency of mutations to cluster on the same chromatid (20-22), and the mechanism and strength of the correlation between mutation rate and recombination rate (23-25). Consequently, distinguishing unexpected clusters of related mutations from genuine non-human fragments is not trivial.

D statistics assume explicitly that mutation rate is constant (18), or at least does not vary enough to distort the expectation of symmetric base-sharing. If this assumption holds, then inter-breeding with archaic hominins offers the only viable mechanism capable of generating asymmetrical base-sharing and significant deviations from zero can be used to help understand patterns human migration (26) and selection (27). However, if the assumption of mutation rate constancy is relaxed, an alternative explanation becomes possible, based on the genome-wide mutation rate being higher among humans who stayed in Africa compared with those who migrated out to colonise the rest of the world (Figure 1). This alternative model can be seen as the converse of the inter-breeding model: instead of non-Africans being unexpectedly *similar* to Neanderthals, Africans would be seen as being unexpectedly *dissimilar*.

**Figure 1.**
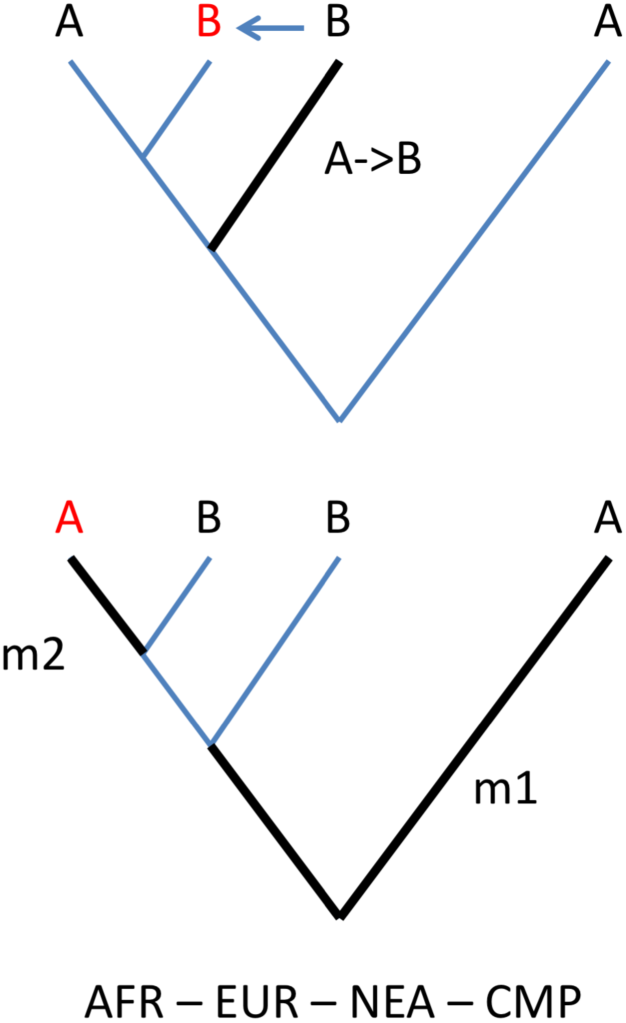
Alternative hypotheses to explain excess base sharing with Neanderthals. Each tree depicts a mechanism by which an “ABBA” pattern is generated, wherein a European human (EUR) shares a base with the Neanderthal (NEA) while, at the same site, an African human (AFR) shares a base with the chimpanzee, Pan troglodytes (CMP). Both trees assume only two states, ‘A’ and ‘B’, with CMP = ‘A’. Heavy black lines indicate lineages on which mutations occurred and key bases appear in red. Top panel: a single mutation creates a new ‘B’ allele in NEA alone which then enters EUR by inter-breeding. Bottom panel: a first mutation m1 creates a “BBBA” pattern then a back-mutation m2 then recreates an ‘A’ allele in AFR.

Mutation rate variation occurs within many groups of organisms, including higher primates (28) though the mechanisms remains largely unclear (29, 30). Among human populations there are reports both of differences in the mutation process (31, 32) and of variation in mutation rate, though the latter evidence is inconsistent, one study reporting a higher rate in Africans (8) and a second finding a lower rate (33) (see also discussion below). One possible mechanism capable of driving variation in mutation rate is heterozygote instability (HI) (5, 34), whereby gene conversion events that target heterozygous sites in heteroduplex DNA formed during synapsis (35) provide extra opportunities for mutations (34). Under HI, the loss of heterozygosity as humans migrated out of Africa (36, 37) would have reduced the mutation rate in non-Africans (8). Although not yet widely accepted, the HI hypothesis enjoys support from microsatellite data (5, 38, 39), from the correlation between heterozygosity lost by humans out of Africa and excess mutation rate in Africa (8) and, more recently, from direct mutation counting in parents and their progeny (7). HI is also consistent with patterns of SNP clustering in humans (20).

Here I attempt to address the question of whether any of the observed asymmetrical base-sharing can be attributed to mutation slowdown rather than inter-breeding with archaic hominins. To do this I construct a series of tests designed to yield opposing predictions under the two models, finding, for the most part, that mutation slowdown is better supported. Given that the processes associated with mutation slowdown are poorly understood, I do not attempt to explain the large body of recent observations relating to archaic hybridisation but hope that my observations will stimulate future research in this area.

## Results

### Mutation rate variation in modern humans and heterozygosity

A key requirement of the mutation slowdown hypothesis is that, since humans migrated out of Africa, the mutation rate inside Africa has been appreciably higher than outside. A three taxon D, D(*Europe, Africa, chimpanzee*), has given conflicting results. The first study, based on Complete Genomics data, found a higher mutation rate in Africa (8), while a second used the Simons Genome Diversity panel to show a higher rate outside (33). Repeating the analysis using data from the 1000 genomes project (40), I found a lower rate in Africa, in agreement with Mallick et al. (33). Although it is unclear why my first analysis gave conflicting results, the key quantity in the context of ABBA-BABA is the relative mutation rate in Africans and non-Africans *since* humans left Africa, and this is not necessarily reflected when all variants are considered.

Since population bottlenecks impact greatly on the allele frequency spectrum, it is not valid simply to focus on alleles rare enough mostly to have arisen since humans left Africa. However, the large loss of heterozygosity non-Africans suffered was modulated across the genome by selection (36, 37, 41). Consequently, if HI operates, then changes in heterozygosity will drive parallel changes in D, creating a correlation between heterozygosity and D. Specifically, excess mutation rate in Africa should be greatest in regions of the genome where excess heterozygosity in Africa is greatest. On the other hand, since D is not impacted by demography (8, 33), if HI does not operate then a correlation cannot exist. In practice, heterozygosity difference and D are correlated in a way predicted by HI (illustrated for all pairwise population comparisons involving GBR, Figure 2). The complexity of the profiles yet strong overall similarity of shape between regions argues for a common mechanism.

**Figure 2.**
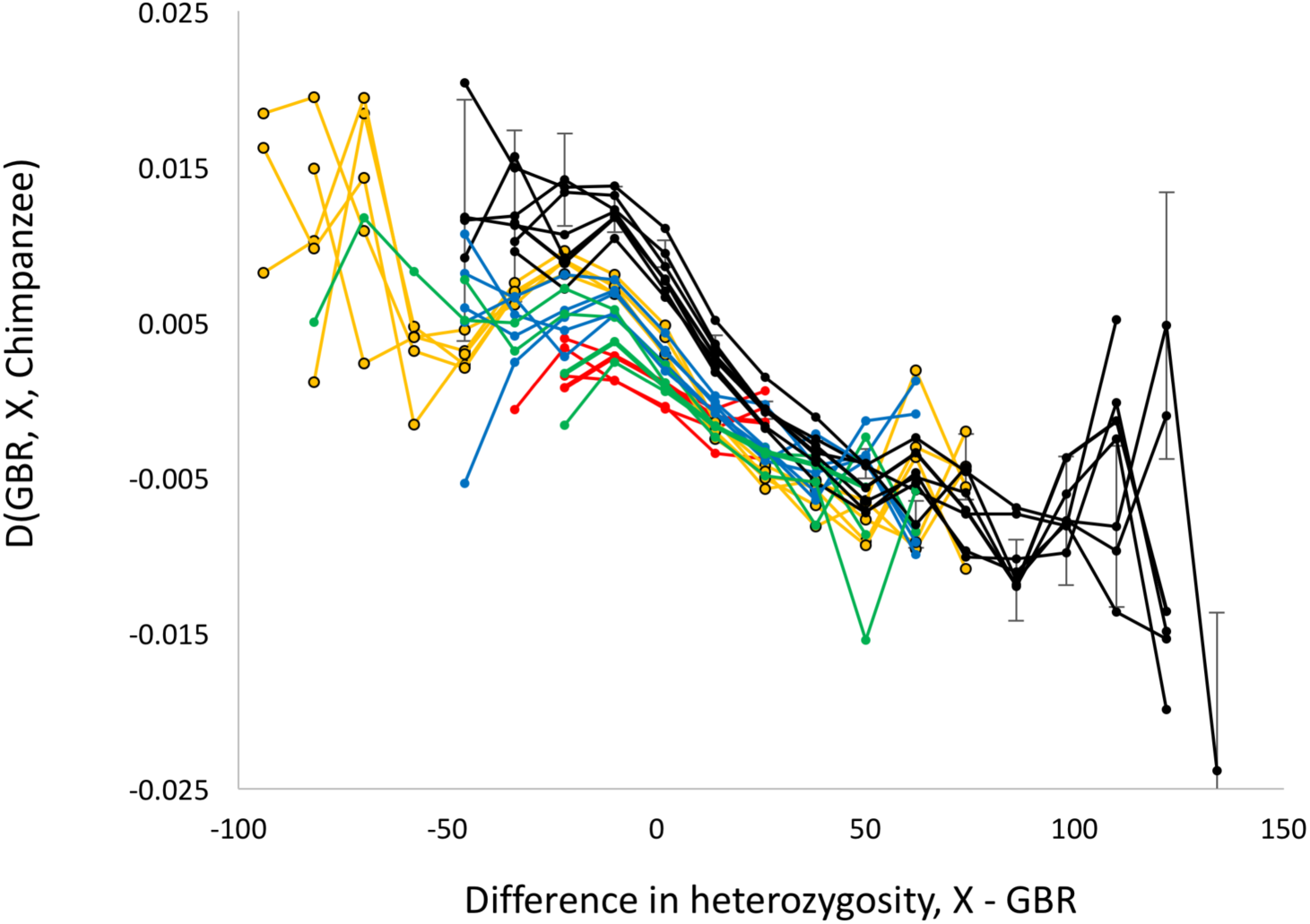
Relationship between heterozygosity difference and relative mutation rate within the genome. Data presented are for all pairwise population comparisons including GBR and a second population, X, colour-coded: Africa, black; Europe, red; South Asia, blue; East Asia, yellow; America, green. To construct this plot, the autosomal genome was divided into 1Mb blocks and each one used to measure the relative mutation rate, expressed as D(GBR, X, chimpanzee) and heterozygosity difference, expressed as the difference in expected number of heterozygous sites. Data were then binned by heterozygosity difference to generate average D values and data for bins with >10 values plotted. For clarity, example error bars, 1 standard error of the mean, are only included for one African population. D is calculated as (BAA – ABA)/(BAA + ABA) such that +ve D values indicate a higher mutation rate in GBR relative to X.

I next tested more directly whether recent mutations occur preferentially in regions of elevated heterozygosity. Even though mutation hotspots are known to evolve rapidly (42), they should be similar in strength and locations among human populations drawn from the same geographic region. Nonetheless, drift will lead to variation in heterozygosity between populations and across the genome. If HI operates, new mutations should be more frequent in population / genomic location combinations that by chance carry higher heterozygosity. Detecting genuine de novo mutations requires deep sequencing of many parent-offspring trios. However, variants occurring just twice in the 1000 genomes data (= ‘doubletons’) are usually less than 500 generations old (43), so their distribution can conveniently be used as a surrogate measure of where mutations are currently most likely to occur.

I identified all doubletons in the 1000 genome data where both copies occurred in the same population. For each doubleton I calculated heterozygosity within 1kb either side, both in the population where the doubleton occurred, *HetMut*, and as an average for the same window across each of the other populations from the same geographic region, *HetOther*. In 24 of the 26 1000g populations the average difference between *HetMut* and *HetOther* is highly significantly positive (Table 1). The two exceptions are Finland and Peru, both populations that have the lowest heterozygosity in their region. After correcting for differences in genome-wide heterozygosity, the all populations show similar, highly significant, positive differences (mean = 0.022 +/-0.004 s.d., range = 0.018 to 0.038). Thus, recent variants do seem to occur preferentially in genomic regions where heterozygosity is higher.

**Table 1:** The impact of heterozygosity on mutation probability. *Recent mutations, defined as variants present in only two copies, both in a single population (‘pop’, for full names see methods), were identified and used to define a 2kb window, 1kb either side of the variant. ‘Difference’ is the average of the difference in heterozygosity between the population where the doubleton occurs minus heterozygosity in the same window averaged over all other populations from the same major geographic region. In all but two cases the average difference is massively significantly positive, the two exceptions being FIN (non-significantly negative) and PEL (massively significantly negative). ‘Dif_adj’ is ‘difference’ adjusted for the difference genome-wide heterozygosity across all windows, revealing a very similar positive values of just over 2% in all populations.*

| pop | difference | s.e. | t | diff_adj |
| --- | --- | --- | --- | --- |
| GBR | 0.0131 | 0.0008 | 17.0 | 0.023 |
| FIN | -0.0014 | 0.0009 | -1.6 | 0.019 |
| CEU | 0.0181 | 0.0008 | 23.5 | 0.022 |
| IBS | 0.0388 | 0.0008 | 46.6 | 0.020 |
| TSI | 0.0324 | 0.0006 | 53.3 | 0.018 |
| CHB | 0.0201 | 0.0006 | 31.3 | 0.019 |
| CHS | 0.0133 | 0.0005 | 25.4 | 0.021 |
| CDX | 0.0187 | 0.0005 | 34.7 | 0.023 |
| KHV | 0.0357 | 0.0005 | 69.8 | 0.022 |
| JPT | 0.0153 | 0.0006 | 26.5 | 0.019 |
| STU | 0.0109 | 0.0005 | 21.8 | 0.020 |
| ITU | 0.0205 | 0.0004 | 51.3 | 0.020 |
| PJL | 0.0177 | 0.0005 | 38.9 | 0.019 |
| BEB | 0.0296 | 0.0005 | 56.0 | 0.021 |
| GIH | 0.0271 | 0.0007 | 40.0 | 0.025 |
| LWK | 0.0308 | 0.0004 | 77.5 | 0.022 |
| ESN | 0.0149 | 0.0006 | 24.2 | 0.024 |
| MSL | 0.0538 | 0.0005 | 104.3 | 0.026 |
| ACB | 0.0126 | 0.0007 | 18.5 | 0.021 |
| GWD | 0.0107 | 0.0005 | 23.1 | 0.019 |
| ASW | 0.0183 | 0.0011 | 16.4 | 0.038 |
| YRI | 0.0322 | 0.0006 | 51.5 | 0.024 |
| PUR | 0.0039 | 0.0011 | 3.6 | 0.032 |
| MXL | 0.0967 | 0.0007 | 137.1 | 0.020 |
| CLM | 0.1716 | 0.0009 | 187.7 | 0.019 |
| PEL | -0.1757 | 0.0016 | -111.0 | 0.026 |

### Are there sufficient numbers of back-mutations?

The mutation slowdown hypothesis requires large numbers of back-mutations. Until now, back-mutation have been assumed to be too rare to be important (1, 18), as the following toy example illustrates. If the mutation rate is 10^−8^ per base per generation and 2000 generations have elapsed since ‘out of Africa’, the 8,156,936 BBBA states reported by Green et al. ((1), Supplementary Materials p. 138) would yield only 163 back-mutations. Even if all these back-mutations occur in Africa, the number is still 50 fold fewer than the observed ABBA-BABA excess of around 8,000. However, this calculation is naïve because real mutations are strongly clustered (21, 22). Since the probability of a back-mutation scales with mutation rate squared, back-mutations are disproportionately likely in mutation hotspots, and likelier still if HI involves the active ‘correction’ of heterozygous sites by gene conversion (19, 35, 44). Although the true rate of back-mutations is difficult to estimate, several lines of evidence suggest it may be of the order required by the mutation slowdown hypothesis. Thus, Green et al. list just over 6000 instances each of BCBA and CBBA ((1), SOM p. 138). Both these base conformations require two independent mutations at the same site, at least one of which must be a transversion. Since transitions outnumber transversions by around two to one, despite representing only half the possible changes, the relative probability of a transition to a transversion is around four (45), implying around 60,000 ABBAs and BABAs due to back-mutation in the Green et al. dataset and 240,000 genomewide (Green et al. generate alignments for about one quarter of the genome). Even this number is likely to be an under-estimate. First, sites with three plus states are probably under-reported, particularly in lower coverage genomes. Second, site-specific and sequence-context considerations (45-47) may cause some sites to mutate mainly by transitions while others would favour transversions. Third, if heterozygous sites are recognised and ‘repaired’ by gene conversion as they are in yeast (35), this would provide a directed mechanism that actively promotes back-mutations.

### Consequence of Excluding Neanderthal sequences

Mutation slowdown and inter-breeding give opposing predictions concerning the dependence of D on the Neanderthal genome. In the inter-breeding model, D will show complete dependence on inclusion of Neanderthal sequences, because Neanderthal bases dictate which sites are informative. Conversely, under the mutation slowdown model, asymmetrical ABBA BABA counts are endogenous to modern humans so different outgroups will give similar results. To test this prediction I used GBR as a reference population and compared D(X*, GBR, Neanderthal, Chimpanzee*) with D(X*, GBR, AA, Chimpanzee*), where X is one of 25 non-GBR 1000g populations and ‘AA’ is the ancestral human allele inferred by 1000g. To prevent any possible spill-over signal, all informative Neanderthal sites contributing to D(X*, GBR, Neanderthal, chimpanzee*) were excluded from the calculation of D(*X, GBR, AA, chimpanzee*). The resulting patterns are highly similar and correlated with each other (Figures 3a, 3b), suggesting a common cause.

**Figure 3.**
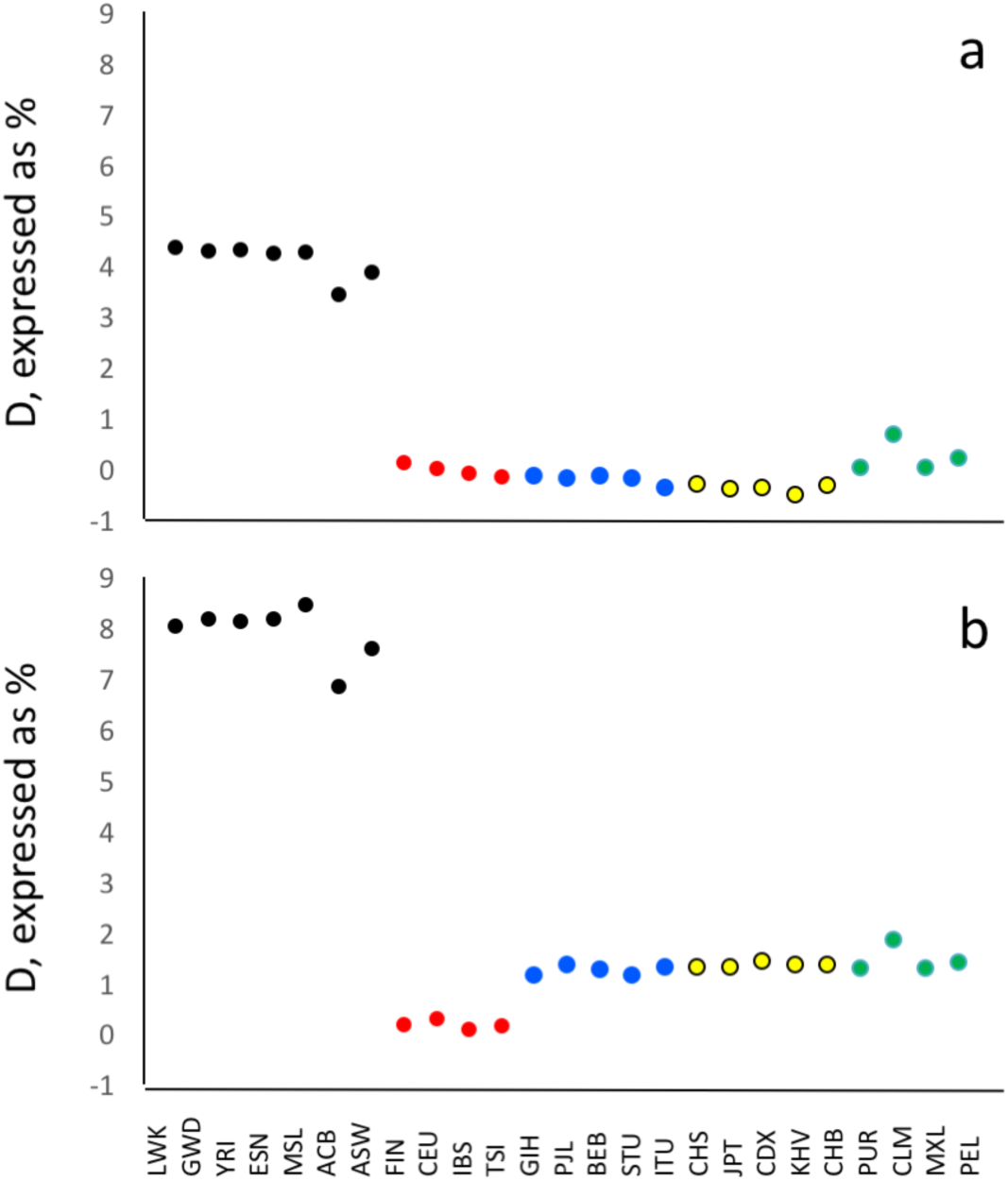
Dependence of D on the Neanderthal genome. Each dot represents an autosomal ABBA-BABA test, expressed as D(X, GBR, Y, chimpanzee), where X is one of 25 other 1000g populations, Y is either Neanderthal (panel 3a) or the inferred ancestral human allele (Panel 3b). Populations are colour-coded by geographic region as in Figure 2, with populations appearing in the same order as listed in methods. Standard errors of the mean are of the order 0.001 and are too small to show. All informative sites contributing to D in 3a were excluded from the analysis in 3b.

Interestingly, D actually strengthens when all Neanderthal information is removed. This strengthening might be expected. When the ancestral allele is used as a form of outgroup, the time-depth of the hominin trio is shallower than when the Neanderthal is used. Consequently, the asymmetrically distributed ABBA-BABA counts will make up a relatively larger proportion of all ABBA-BABA states. The fact that larger Ds result when Neanderthal information is excluded indicates that, in this analysis, Neanderthal introgression cannot be the dominant mechanism driving positive D. This point is emphasised by the fact that replacing Neanderthal and chimpanzee with various combinations of higher primates (gorilla and orangutan) also generate significant D (data not shown).

### Which allele is rare?

Inter-breeding and mutation slowdown also give opposing predictions for the frequencies of the ‘A’ and ‘B’ alleles. Under inter-breeding, positive D is driven by Neanderthal ‘B’ alleles entering human populations outside Africa. Under mutation slowdown, positive D is driven by an excess of ’A’ alleles generated by back-mutation in Africa. In both cases, the key alleles will tend be rare, having entered their respective populations after humans left Africa. However, the patterns are opposing. Inter-breeding and mutation slowdown predict that the ‘B’ allele in Europe, will be rare and common respectively. To test this prediction I partitioned autosomal D(*X, GBR, Neanderthal, chimpanzee*) by the frequency of the ‘B’ allele in GBR. When *X* is African, D averages 75.7%, -59.1% and -0.05% depending on whether the ‘B’ allele is at high (>90%), low (<10%) or intermediate frequency respectively (Figure 4). Since positive D occurs when ‘B’ is common in GBR, these results support mutation slowdown over inter-breeding. The highly polarised patterns seen when one allele is rare / common and virtual lack of any signal at intermediate frequencies (mean D = 0.05%, Figure 4c) argues against mechanisms where appreciable numbers of alleles are at intermediate frequency such as very ancient inter-breeding.

**Figure 4.**
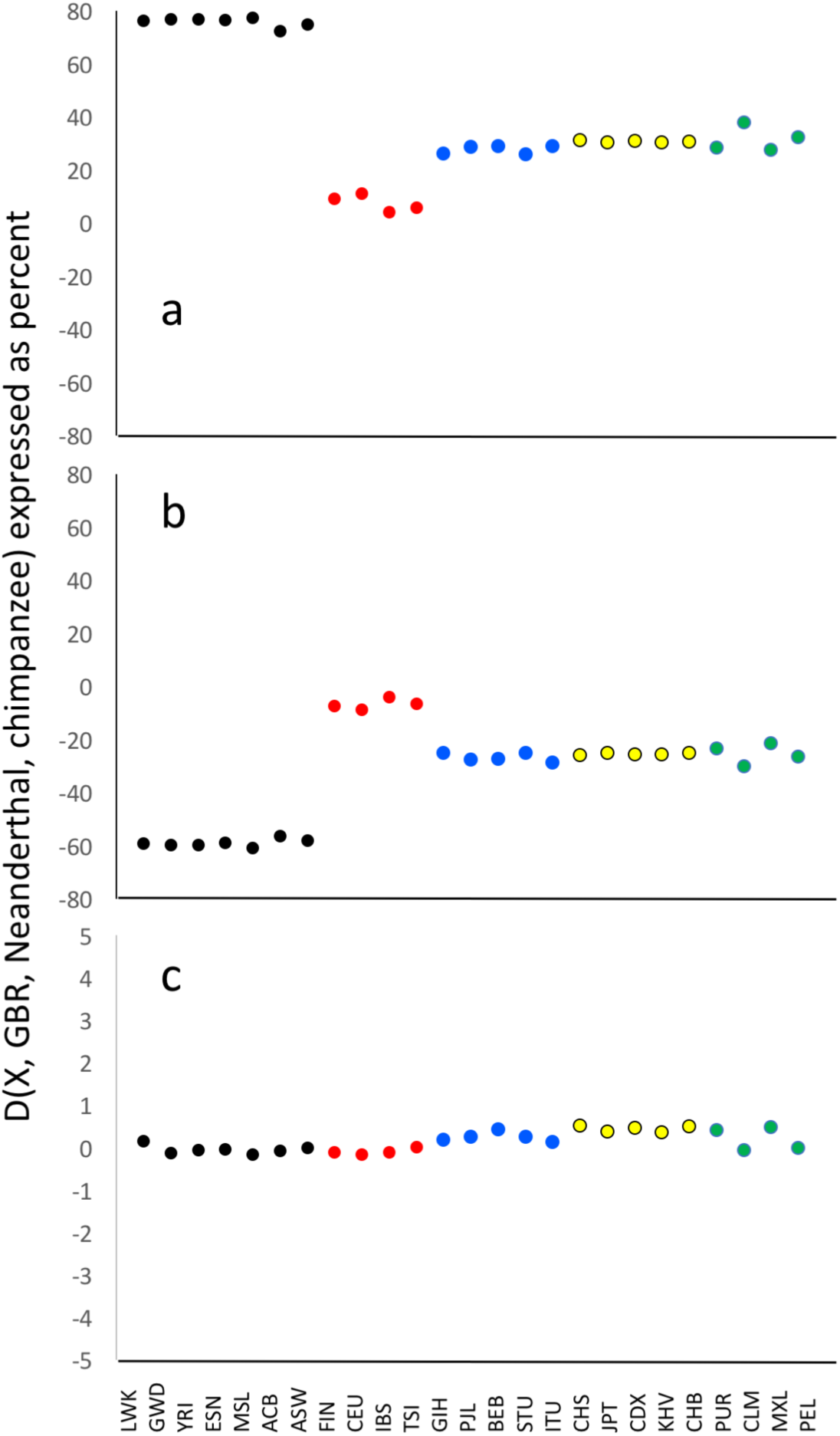
Dependence of D on the frequency of the ‘B’ allele. Data are subsets of those in Figure 3, see that legend for details. The data are partitioned according to whether the frequency of the ‘B’ allele in GBR is common (>90%), rare (<10%) or intermediate frequency, shown in Figures 4a, 4b and 4c respectively. Note how the classic observation that D(AFR, EUR, NEA, CMP) ∼4% reflects an average of the extreme patterns seen in 4a and 4b. NB the vastly different scales on the Y axis.

### Which populations drive most variation in D?

A further feature that could help distinguish the two hypotheses relates to variation in D among different population combinations. Both hypotheses predict one region drives D while the other acts as a passive control. Consequently, across all African – non-African population combinations, rotating populations from one region should cause more variation in D than rotating populations from the other. Specifically, D should vary more when non-African and African populations are rotated under the inter-breeding mutation slowdown hypotheses respectively. Ignoring the highly admixed ‘American’ populations, variance in D is appreciably greater when African populations are rotated against non-African populations (mean variance = 3.33×10^−4^, +/-1×10^−6^ s.e.m.) than *vice versa* (mean variance = 2.54×10^−4^, +/-2.9×10^−6^ s.e.m.), see Figure 5. If anything, this lends further support to the mutation slowdown hypothesis.

**Figure 5.**
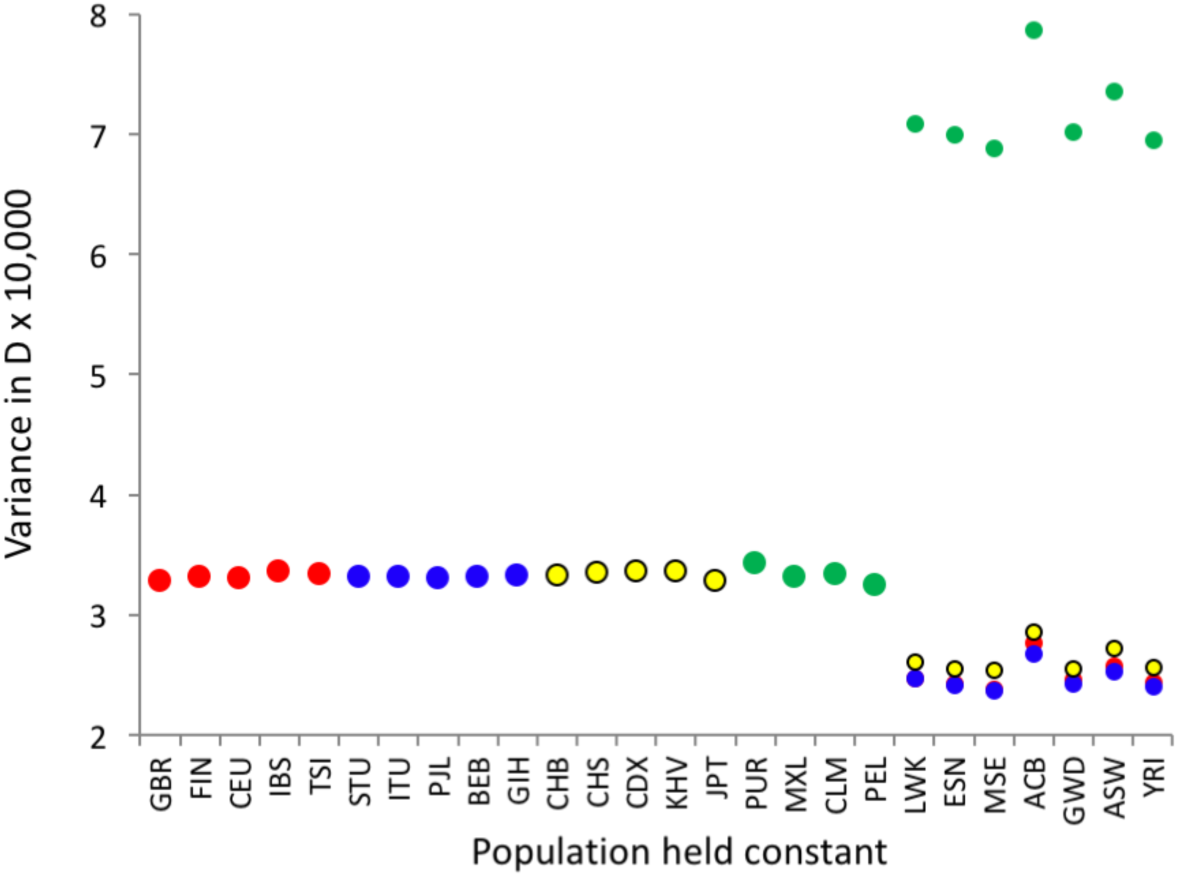
Dependence of the variance in D on which populations are rotated for African – non-African population pairs. D(H1, H2, Neanderthal, chimpanzee) values were generated for all African – non-African population pairs, H1, H2. For each population (X-axis) variance in D (Y-axis) is calculated for all comparisons in which it is included. Non-African variances are based on all comparisons to the African populations (N=7) while the African comparisons are partitioned by each of the other four population groups (Europe, N=5; Central Southern Asia, N=5; East Asia, N=5; America, N=4). Colour coding of non-African populations follows Figure 2.

By far the highest variance is seen in African populations when American populations are rotated. Since the variances are so different from any of the other regions it seems likely that these extreme values relate to the high but varying levels of admixture. Arguments could be made in support of both hypotheses. Thus, the high variation in D could be driven by increased variation in the frequency of introgressed fragments. Equally, if mutation slowdown is driven by the loss of heterozygosity that occurred ‘out of Africa’, highly admixed populations would likely exhibit a counter-trend. Distinguishing these two possibilities is not trivial and has not been attempted here.

### Resolving the relationship between D and heterozygosity

Several studies have sought to understand how introgressed Neanderthal genes have been impacted by natural selection (2, 48). The action of selection can be captured by B-statistics, a derived measure of the extent to which patterns of heterozygosity have been distorted by selection (49). B and D are positively correlated across the genome, interpreted as selection modulating the frequency of introgressed fragments (50). However, a correlation between D and heterozygosity is also expected under mutation slowdown. Here, the amount of diversity lost in the out of Africa bottleneck was modulated by natural selection, acting to accelerate loss of diversity in some genes and to reduce loss in others (51). As a result, strongly selected genes such as those in the immune system may have experienced more or less mutational slowdown, depending on whether they came under directional or balancing selection respectively. In turn, this could drive unusually high or low D values around selected genes.

Although both hypotheses predict a relationship between D and heterozygosity, the exact nature of the relationship differs. Inter-breeding predicts a more or less simple relationship between heterozygosity and D in non-Africans only. In contrast, for mutation slowdown the key quantity is *difference* in heterozygosity and, while the main focus is on African – non-African comparisons, HI should drive non-zero D wherever heterozygosity differences have persisted for appreciable amounts of time. A second important difference is that, while selection acting on Neanderthal fragments may drive a correlation across regions within a genome, its impact on genome-wide values for individuals or populations should a minimal. Moreover, if Neanderthal fragments are common enough to increase heterozygosity, their presence would drive a negative relationship between heterozygosity difference and D. In contrast, HI should drive a positive correlation both within *and* between genomes.

To explore the impact of genome-wide heterozygosity difference on D(*H1, H2, Neanderthal, chimpanzee*), I considered all pairwise population comparisons in the 1000g data. African – non-African combinations show high D and large differences in heterozygosity, so tell us little about the likely mechanism. I therefore focused on comparisons either inside or outside Africa, where differences in Neanderthal content are expected to be minimal or negligible (Africa). As expected, D values are small, rarely exceeding 1%. Nonetheless, outside Africa D is strongly predicted by difference in heterozygosity (black symbols, Figure 6), with an r^2^ of 81.2%. Since there is no *a priori* reason why a given population should appear as *H1* rather than *H2,* each comparison is included twice. This removes possible bias due to which population is made H1 but means that the r^2^ value cannot be interpreted directly. Within Africa, non-admixed comparisons (red) conform quite closely to the outside Africa trend. In contrast, comparisons involving the two admixed populations (ACB, ASW, in blue) show a similar slope but much larger D, reinforcing the idea that admixture impacts D more than expected from passive mixing.

**Figure 6.**
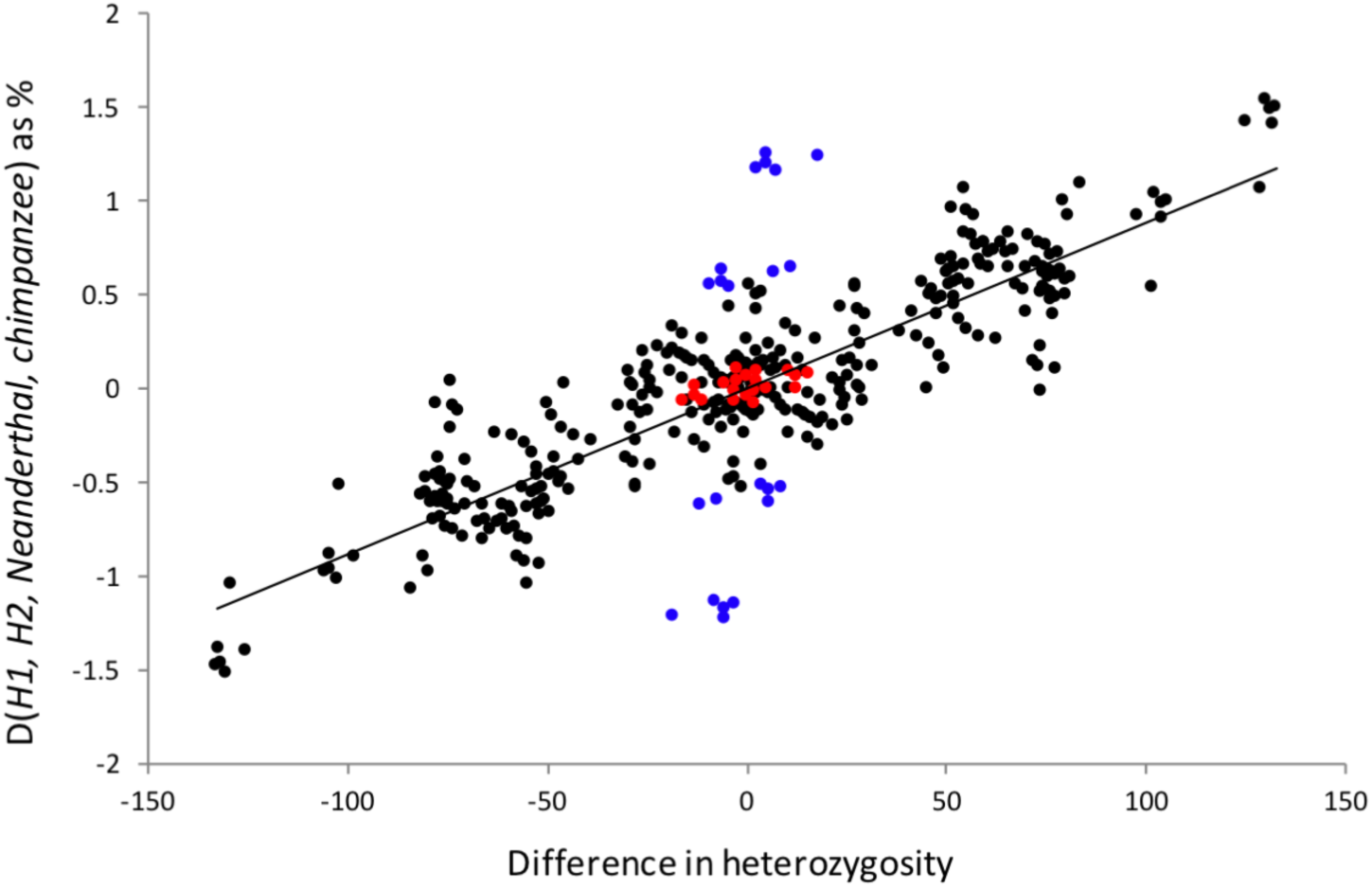
Heterozygosity difference predicts D in pairwise population comparisons. Autosomal D(H1, H2, Neanderthal, chimpanzee) was calculated for all pairwise combinations of non-African populations (N=19 populations, black dots), non-admixed African population pairs (N=5 populations, red dots) and within Africa comparisons involving either admixed population, ACB or ASW. The X-axis is difference in mean number of heterozygous sites per megabase between populations, expressed as H1 - H2. The Y-axis is D which, being generally small, has been expressed in percent. The positive slope indicates that when H1 has higher heterozygosity, population H2 carries more bases in common with the Neanderthal. Note, each population pair is represented twice, with H1 and H2 reversed, making the pattern symmetrical about zero.

A next examined the impact of heterozygosity difference on D across regions *within* the genome, I divided the autosomal genome into 1Mb windows. Within each window and for each of all possible pairwise population combinations I calculated heterozygosity in each population (Het_H1_ and Het_H2_) and counted ABBAs and BABAs for the tetrad *H1, H2, Neanderthal, chimpanzee*. I then fitted multiple regressions of the form

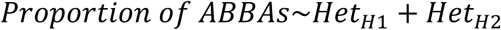

where the proportion of ABBA counts is fitted as a binomial response. 78 windows excluded where the total number of ABBAs + BABAs was less than one (fewer than 3% of windows, in effectively all cases windows with no chimpanzee alignment or including big overlaps with centromeres / telomeres). Apart from 20 non-significant, within Africa comparisons, the slopes of H1 and H2 are invariably significant (P<0.01, though for a large majority P<<10^−10^) and show opposing slopes, being positive and negative respectively (Figure 7).

**Figure 7.**
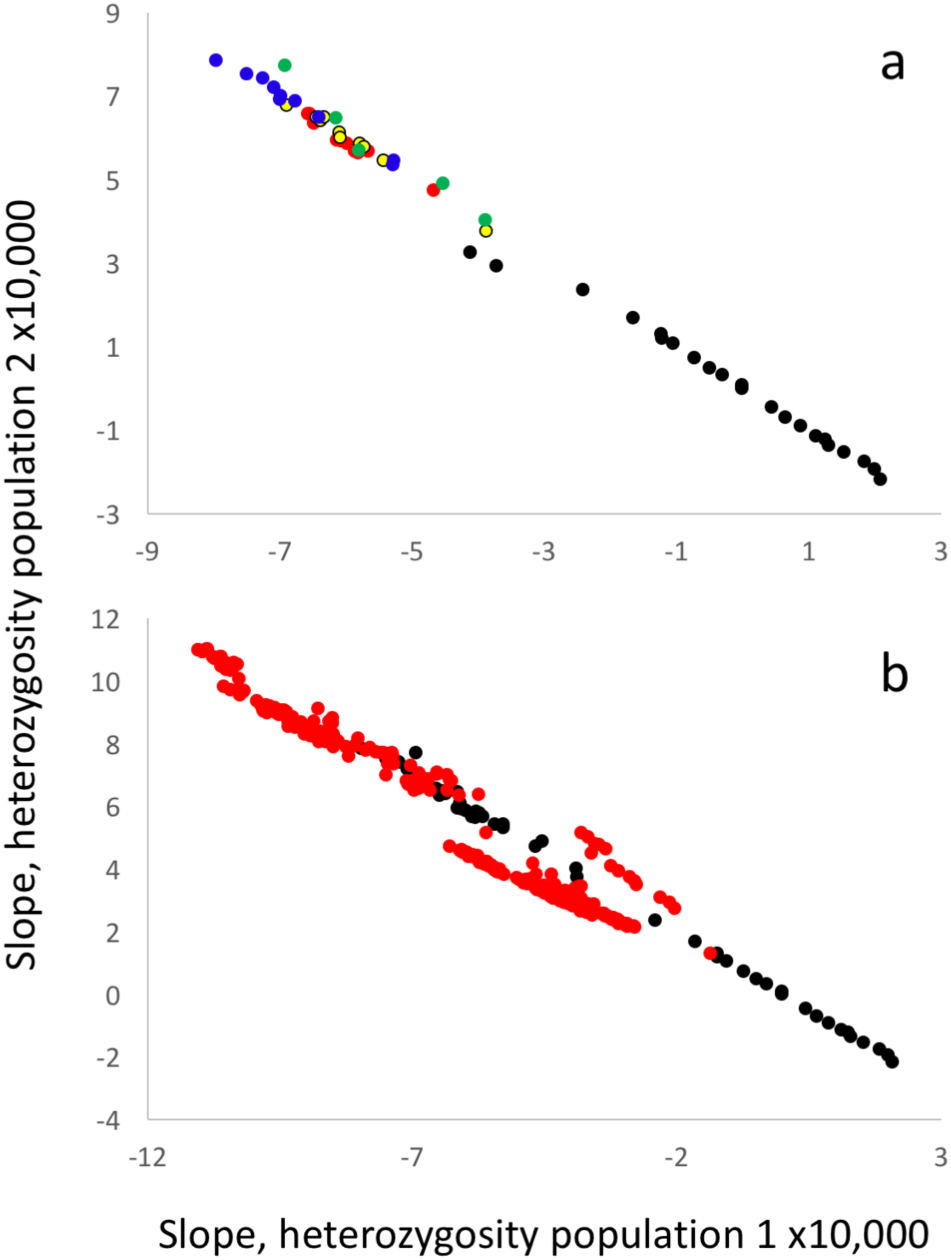
Relationship between heterozygosity and D across the genome. The autosomal genome was analysed in non-overlapping 1Mb windows in all pairwise combination of the 26 1000g populations. For each window in each pair I counted the numbers of ABBAs and BABAs and the expected number of heterozygous sites in each population (Het1, Het2). I then fitted a binomial general linear models of the form ABBA/BABA ∼ Het1 + Het2. Slope estimates for all within-region population pairs are shown in Figure 7a, colour coded as in Figure 2. Figure 7b depicts the full set of comparisons, coded within region = black, between regions = red. All slopes are significant except for 20 within-Africa comparisons (righthandmost points in Figure 7a).

Since previous studies only consider a single heterozygosity, expressed as a B statistic (49, 50), I was interested to see if the two heterozygosity model (2HM) had similar or better explanatory power. For each population combination I compared the better fitting (lower AIC) single heterozygosity model (SHM) with the 2HM. No difference was found within Africa (N=21 comparisons, mean AIC difference 0.1). In all other instances, the 2HM AIC was substantially lower (Table 2a). Apart from within Africa, the 2HM explained 2-9.5% of variation in D, 3-112 times more than the best SHM (Table 2b). Combined, the massive superiority of the 2HM and the highly significant, always opposing slopes are difficult to reconcile with a model based on selection, yet support a model where the key quantity is *difference* in heterozygosity.

**Table 2:**
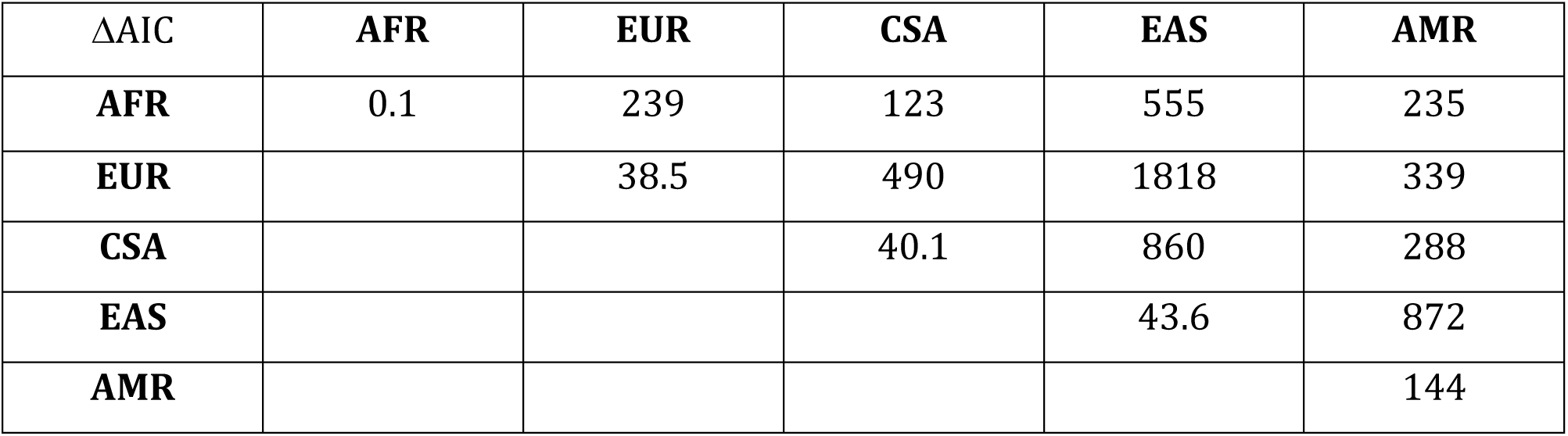
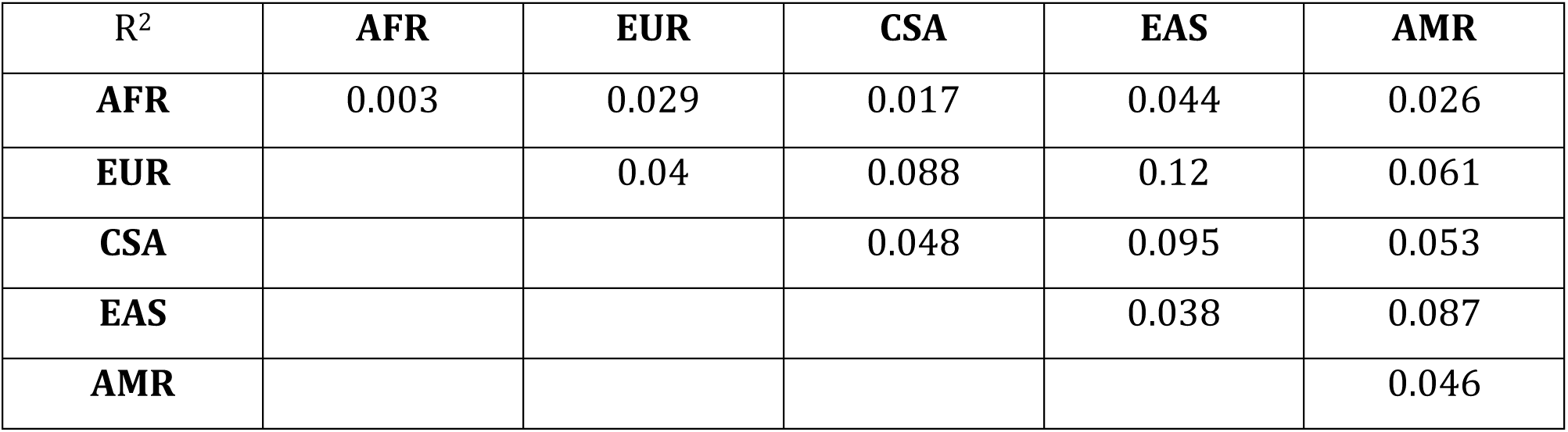
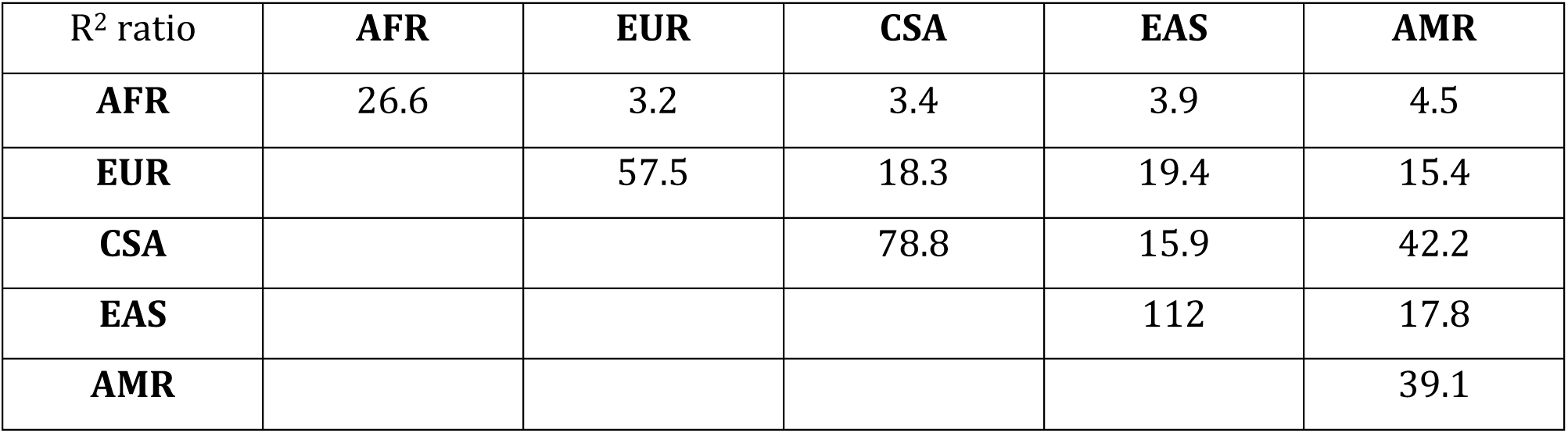
Relationship between heterozygosity and D. *For each possible pairwise population comparison ABBA, BABA and heterozygosity counts were made for all 1Mb autosomal windows. General linear models were fitted with binomial response ABBA and BABA counts and predictors either both population heterozygosities (2HM) or just one (SHM). Each 2HM was compared with the better fitting SHM and summary statistics averaged by regional comparison: for Africa (AFR), Europe (EUR), Central Southern Asia (CSA), East Asia (EAS) and America (AMR): Table 2a is difference in AIC; Table 2b is proportion of variance in D explained by the 2HM; Table 2c gives the ratio of variance explained by the 2HM divided by variance explained by the best SHM.*

### Do non-Africans carry unusual fragments?

D is a relative measure and does not distinguish between non-Africans being more similar to Neanderthals and Africans being less similar. For a more objective estimate of the frequency of putative introgressed fragments I calculated all-against-all intra-population pairwise divergences for non-overlapping 20Kb windows across the genome, recording the maximum value in each population (MaxPD). With generally higher diversity, MaxPD(Africa) should be higher than MaxPD(outside Africa). However, even a single Neanderthal (or other archaic) fragment in a non-African population will tend to reverse this expectation. In other words, the ratio MaxPD(outside Africa)/MaxPD(Africa) should be <1 when introgressed fragments are absent and >1 when present. Since previous studies suggest that very few genomic regions carry zero Neanderthal ancestry (11, 50), the inter-breeding hypothesis predicts a ratio >1 in most windows.

Only ∼5% of autosomal windows show a ratio >1 (exemplified by chromosome 1, Figure 8a). Moreover, since a typical Neanderthal sequence is more divergent from modern humans than any two modern humans are from each other, genuine Neanderthal fragments should often yield even higher ratios. Setting the threshold ratio to 1.25 and 1.5 sees the proportion of all windows that qualify as putatively archaic fall to 0.3% and 0.08% respectively. Importantly, the locations of the few large ratios that are present do not correlate with peaks in the published Neanderthal landscape (Figure 8b) (50). Also, introgression has previously been inferred to be near-zero on the X chromosome (50). If high ratios do indicate introgressed fragments they should be rarer on the X. In fact, if anything, the X-chromosome carries more windows with ratios >1 than then autosomes (Figure 8c).

**Figure 8.**
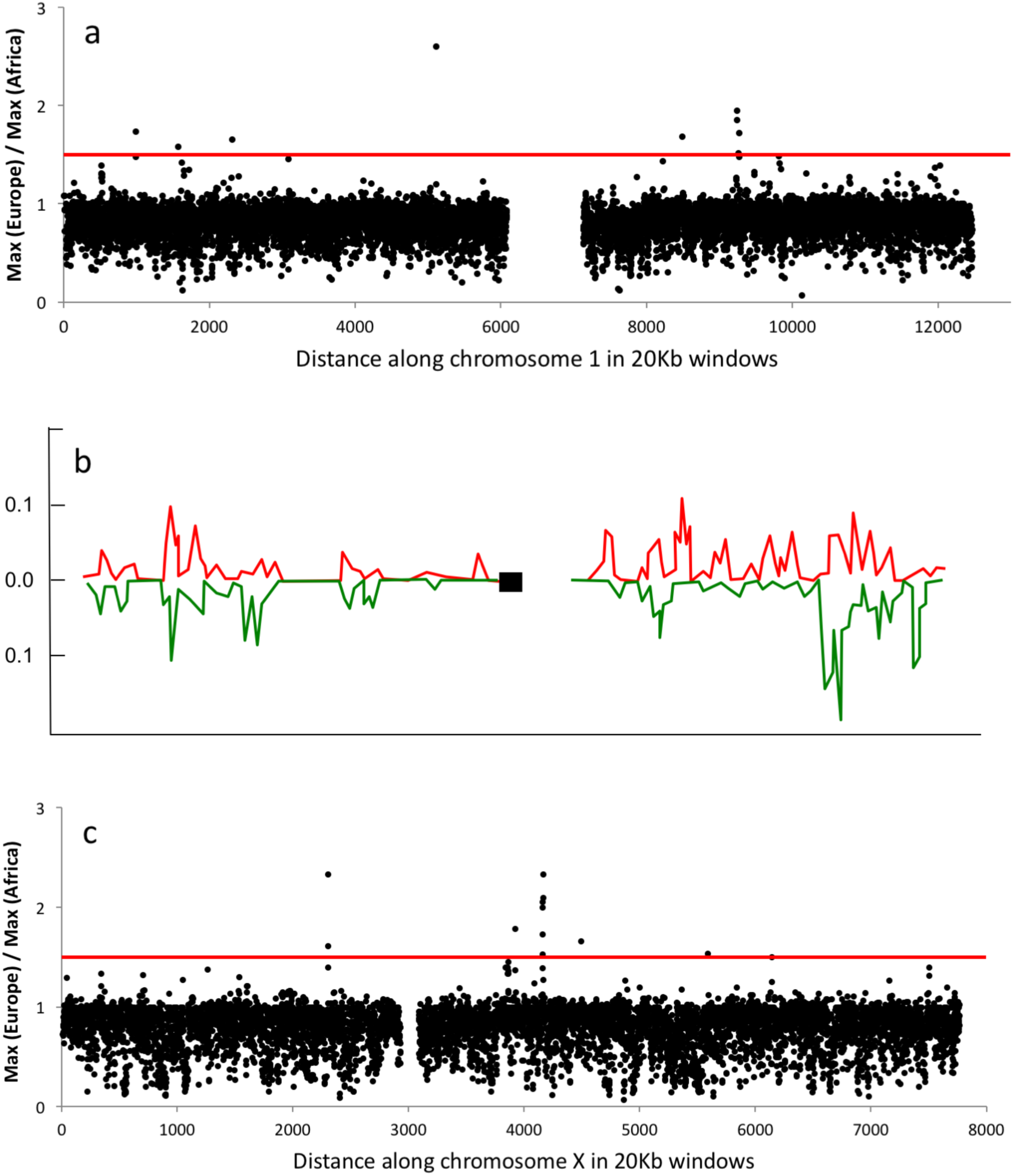
A general test for presence of introgressed archaic fragments outside Africa. Figure 8a shows the ratio of maximum sequence divergence in Europe (GBR, FIN, CEU, TSI, IBS) to maximum sequence divergence in African (LWK, ESN, MDL, GWD, YRI) for 20Kb windows on chromosome 1. All other autosomes look very similar. Almost all windows give ratios less than the 1.5 expected if a Neanderthal fragment was present in one of the European samples (red line), and the average ratio is ∼0.8, reflecting the greater diversity in Africa. For comparison, Figure 8b gives the landscape of inferred Neanderthal contribution to Europeans (red) and East Asians (green), redrawn from Sankararaman et al. (50). Published studies suggest little Neanderthal contribution to the X chromosome. If the few high ratios reflect introgressed fragments they should be rarer on the X but appear to be just as common (Figure 8c).

Even though many fewer windows yield large MaxPD(outside Africa) compared with the published Neanderthal landscape (50), where regions lacking introgressed fragments appear rare, the high ratios could still reflect genuine archaic sequences. If so, MaxPD(outside Africa) and D should be positively correlated: high ratios should occur in windows that also contribute large D. Plotting D against MaxPD(Europe) for all autosomal 100kb windows reveals a highly significant *negative* correlation, with highest D being associated with the smallest MaxPD values and the highest MaxPD giving D’s close to zero (Figure 9). Such a pattern is difficult to reconcile with a model where high D values are driven by divergent archaic fragments but is consistent with a model where the highest D values occur in regions where most diversity has been lost ‘out of Africa’.

**Figure 9.**
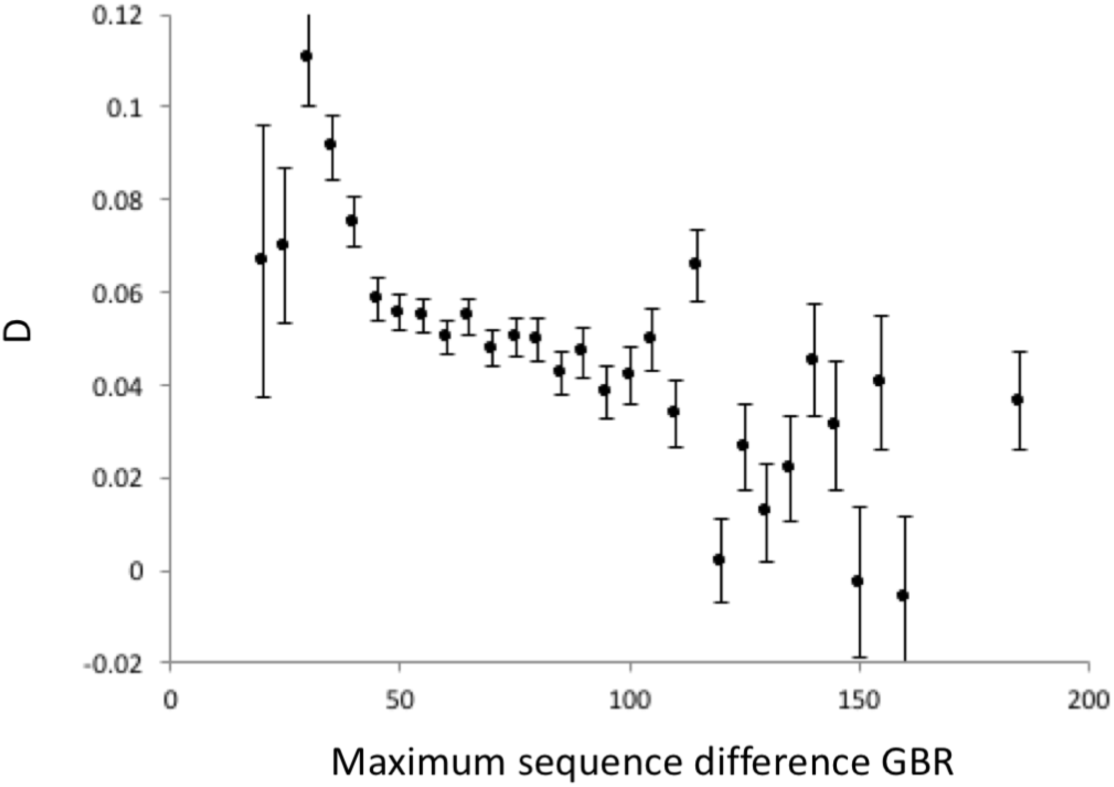
Objectively unusual fragments in non-African populations are associated with low D. For all autosomal data I calculated D(YRI, GBR, Neanderthal, chimpanzee) for each 100Kb window and plotted the resulting values, binned by maximum divergence among fragments within GBR. The largest D values used to infer introgression are strongly associated with low Max(GBR), the opposite of what would be expected if large ratios genuinely reflect the presence of Neanderthal fragments. Windows with the very lowest Max(GBR) do not give the highest D values, plausibly because these include appreciable numbers of sequences that have very low diversity in all populations: these will have low D-values that depress the average.

Finally, I attempted to estimate the proportion of unusual, presumed archaic origin fragments in individual non-African genomes. Within each window I define ‘unusual fragments’ as non-African sequences that exceed MaxPD(Africa) when compared with at least 10 intra-population comparisons. This requirement reflects the rarity of most archaic fragments and also reduces the impact of comparisons between the most divergent non-African human fragments. Since Neanderthal fragments may occur in Africa, I take MaxPD(Africa) conservatively to be the smallest of the five individual African population maxima (ASW and ACB are not included because they have varying levels of European admixture). As an internal calibration, each non-African population was ‘spiked’ with a randomly selected African individual from GWD. In 33% of 20Kb windows this African sample qualified as ‘unusual’. Since Neanderthal fragments will be appreciably more divergent it is reasonable to assume that upwards of 50% of archaic fragments would be detected. Using the 50% value, the average archaic proportion in non-Africans is 0.14%, about 10 fold lower than currently estimated. Moreover, since divergent sequences but can arise through chance, selection and population mixing, even this figure should be treated as an upper limit.

## Discussion

Neanderthals share more nucleotide bases with modern non-African humans than with Africans. This basic pattern can be explained either by historical inter-breeding or by the mutation rate having slowed in humans who left Africa. Here I test a series of opposing predictions aimed at distinguishing inter-breeding from mutation slowdown. By and large, the patterns I uncover favour the mutation slowdown model.

Mutation slowdown requires that mutations rates differ appreciably among human populations and that, since humans migrated out of Africa, the rate has been higher in Africa. Current evidence is inconclusive. Differences between Africans and non-Africans have been reported in mutation rate (8, 33), mutation type (31) and recombination rate (52), though the largest study finds a higher mutation rate outside Africa (33). On the other hand, the latter study does not distinguish between mutations that predate and postdate ‘out of Africa’. When I analyse Complete Genomics data and 1000 genomes data I find significant but opposing trends, perhaps suggesting high sensitivity of the result to data filtering and or levels of imputation.

The question of whether Africans and non-Africans differ in their genome-wide mutation rates *since* humans left Africa thus remains unresolved. However, when mutation rate difference is estimated using D(*African, non-African, chimpanzee*) (33) across the genome, there is a striking correlation with the amount of diversity lost: the more diversity was lost, the greater the excess mutation rate in Africa. Since the dominant mechanism driving heterozygosity difference is drift during the out of Africa bottleneck (37, 41), any causal relationship must be in the direction of changes in heterozygosity influencing mutation rate rather than *vice versa*. Having said this, the data do not formally rule out other, as yet unknown mechanisms. Such mechanisms are difficult to conceive because the most obvious way to change mutation rate would be through a variant polymerase, but this would only impact the genome-wide average, it would not drive a correlation between diversity lost and mutation rate.

A second important issue is the extent to which the mutation slowdown model can explain a raft of observations that support the inter-breeding hypothesis. That some inter-breeding occurred seems likely, based on discoveries of a genuinely hybrid skeleton and introgressed haplotypes (27, 53). However, the case for frequent mating leading to a larger legacy depends on genome-wide patterns which, though reported in many different forms (2, 12, 18, 54), generally fail to distinguish between non-Africans being more similar to Neanderthals and Africans being less similar. Thus, while current evidence for inter-breeding appears compelling, until the various analyses are repeated in a way that distinguishes inter-breeding from mutation slowdown, the conclusion that inter-breeding offers the only possible explanation seems to me to be premature.

It is of course desirable to assess the extent to which the panoply of recent observations currently interpreted as evidence of inter-breeding could be explained by a model based on mutation slowdown. If HI operates, it would drive variation in D (and related measures) wherever heterozygosity varies, either between populations, due to changes in population size (37), or across the genome, due to the action of selection (55). Moreover, recombination rate and mutation rate are correlated (56) and mutations occur non-independently to form clusters (20-22, 57). By interacting with HI, these phenomena appear capable of creating most of the patterns currently taken to indicate inter-breeding. Unfortunately, it is not yet possible to make a proper assessment of the extent to which this potential is fulfilled, if at all, because most of the component processes, although likely present, are too poorly understood to model effectively. It thus seems premature to claim either that HI can or cannot explain a given pattern.

A key exception to the rule that most analyses fail formally to exclude HI appears in the original Green et al. study (1) and exploits the mosaic of sequences with European and African ancestry found in the human reference sequence. A small number of European but not African sequences show high similarity to Neanderthals yet low similarity to the Venter sequence (their Figures 5 and S39). This contradicts my observation that objectively unusual fragments are very rare. Where the truth lies remains to be determined. However, Green et al.’s analyses include multiple stages any one of which could be subject to a filtering bias of the sort that has changed interpretation elsewhere (doi:10.1038/nature.2016.19258). There is also an issue with the raw data. The length of the Neanderthal branch should be less than 10% of the total hominin-chimpanzee divergence but Green at al.’s data suggest an implausibly high figure of ∼75% (see their Table S51: counts of 5,827,247 and 8,156,936 for states AABA and BBBA respectively). In contrast, my analyses are based entirely on high quality (if low coverage) modern sequences, and include an internal calibration in the form of an African individual added to all non-African samples. That the African sample is identified as unusual in a third of all windows suggests that my approach should be capable of detecting ancient fragments, if present, in upwards of 50% of windows.

Despite the above uncertainties, my analyses do uncover a number of additional patterns that appear difficult to reconcile with a model based entirely on inter-breeding. First, D actually strengthens when the Neanderthal genome is replaced by the inferred human ancestral bases. Interestingly, this observation agrees with data presented by Green et al., who report 756,324:689,594 BAAA:ABAA, a ratio of 1.097, similar to but larger than the 1.087 ratio seen in the same Table S51 for BABA:ABBA, and involving seven times as many bases. Green et al. ascribe their 66,730 excess BAAAs to sequencing errors, though this number seems implausibly large for high coverage sequences and it is unclear why sequencing errors would be so asymmetrically distributed. A more parsimonious explanation is that both the similarity of the BABA:ABBA and BAAA:ABAA ratios and the similarity between D(*H1, H2, Neanderthal, chimpanzee*) and D(*H1, H2, ancestral human, chimpanzee*) reflect a common phenomenon, one that is inherent to all comparisons among modern humans and has little or no dependence on which outgroup is used as long as it lies close to the base of modern humans.

A second problem for the inter-breeding story relates to which alleles are common and which are rare. Most early studies focused on individual high coverage genomes so were unable to consider allele frequencies. By using the 1000 genomes data I have been able to show that D is acutely sensitive to whether the ‘B’ allele is common or rare, only becoming positive when the ‘B’ allele frequency exceeds 90% in Europe. This is the exact opposite of what is expected under the inter-breeding hypothesis. Most Neanderthal fragments are likely neutral or near-neutral (55) and would have entered modern humans around 2,000 generations ago, too recently for any but a handful to drift to high frequency. If D is driven mainly by inter-breeding, positive D value should be associated with sites where the ‘B’ allele is rare in Europe, not common as I report. This analysis emphasises the extra information gained by considering allele frequencies.

The third problem involves the relationship between heterozygosity and D, as discussed above. Variation in genome-wide heterozygosity is dominated by the out of Africa bottleneck (37, 41) and should be little impacted by a 1-2% Neanderthal legacy (if present). However, genome-wide D is almost perfectly predicted by heterozygosity difference across all population comparisons, the population with relatively lower heterozygosity invariably appearing closer to Neanderthals. If Neanderthal fragments did have an appreciable effect on heterozygosity this would, if anything, drive the opposite trend, because higher D would be linked to higher heterozygosity driven by introgression. The ubiquitous converse trend both within and outside Africa is therefore at odds with a model based entirely on inter-breeding.

The relationship between heterozygosity and D *within* a genome also argues against inter-breeding. Previous work reveals a correlation between B statistics and D, interpreted as indicating selection acting on introgressed fragments. However, these studies only consider a single heterozygosity. When I fit models that include heterozygosity in each population separately (two heterozygosity models, 2HMs) a dramatically different picture emerges. All pairwise population comparisons, apart from those within Africa, yield a vastly superior fit for the 2HM, and the two slopes invariably go in opposite directions. Such opposing slopes fit well with a model where the key quantity is *difference* in heterozygosity, but seem at odds with a model based on selection, where all significant regressions should go in the same direction.

An interesting and perhaps telling feature of the 2HMs is the relationship between strength of correlation and genetic distance. In African – non-African comparisons, the 2HMs only explain 3-5 times as much variation in D compared with the best single heterozygosity model. In contrast, among non-African comparisons the single heterozygosity models are often non-significant while the 2HMs are at their strongest, particularly in comparisons between the three Eurasian regions EUR, CSA and EAS. This is difficult to understand using a model based on selection, where the strongest correlations should always be the African – non-African comparisons. A plausible explanation based on HI can be made but needs more work to determine whether it works in practice. Thus, populations within a region show little variation in D across the genome because they are too similar in heterozygosity and have not been separated long enough for many mutations to accumulate. Similarly, African – non-African comparisons show strong D but the correlation with heterozygosity is somewhat degraded due to complicated patterns of mixing and demographic change since humans left Africa. This leaves the strongest signal to form over an intermediate timescale where the best balance lies between developing a strong signal and this signal being degraded by complex and often contrasting demographic histories.

Finally, a number of other observations sit somewhat uncomfortably with the idea of sufficient inter-breeding to leave a 2% legacy. Estimates of introgression are near-zero and zero for the X chromosome and mitochondrial DNA respectively. This pattern could be explained by some combination of unidirectional gene flow, mediated by a preponderance of male Neanderthals mating with female humans, and selection acting against the Neanderthal sex chromosomes. However, both mechanisms are speculative and lack empirical support. Were human men unable or unwilling to defend their wives against (presumed) rape and, if defence was not possible, why did the populations not move apart to reduce contact? It is also unclear that mixed offspring would thrive and be accepted readily by society. Here, the context of the hybrid skeleton (9) may be relevant. People generally use caves either for burials, when many remains would be found, or for living, when no skeletons would usually be found. The finding of just two skeletons is not consistent with either but might reasonably be interpreted as the ostracising or separation of individuals deemed uncomfortably different, implying lower fitness.

In conclusion, I explore an alternative explanation for relatively greater base-sharing between Africans and non-Africans, based on mutation slowdown out of Africa. Although the mutation slowdown model is speculative and its ability to account for patterns used to infer inbreeding are largely untested, this new model does make a number of predictions that appear to be fulfilled. In comparison, the inter-breeding struggles to account for the patterns I find. Clarifying which model fits best across all observations will require a lot more work. I hope that future studies will seek both to determine how well mutation slowdown fits for the many published studies, and to find reasonable explanations for why the interbreeding model fits so poorly in the analyses I have conducted. Future inferences about possible inter-breeding should consider mutation slowdown as a viable alternative explanation that needs to be eliminated before introgression can confidently be inferred.

## Methods

### Data

Modern human sequences were obtained from the 1000 genomes project, Phase 3, and downloaded as composite vcf files. These comprise low coverage genome sequences for 2504 individuals drawn from 26 modern human populations spread across five main geographic regions: Africa (**LWK**, Luhya in Webuye, Kenya; **GWD**, Gambian in Western Division, The Gambia; **YRI**, Yoruba in Ibadan, Nigeria; **ESN**, Esan in Nigeria; **MSL**, Mende in Sierra Leone; ACB, African Caribbean in Barbados; ASW, African Ancestry in Southwest US), Europe (**GBR**, British from England and Scotland; **FIN**, Finnish in Finland; CEU, Utah Residents (CEPH) with Northern and Western Ancestry; **TSI**, Toscani in Italy; **IBS**, Iberian populations in Spain), Central Southern Asia (GIH, Gujarati Indian in Texas; **PJL**, Punjabi in Lahore, Pakistan; **BEB**, Bengali in Bangladesh; STU, Sri Lankan Tamil in the UK; ITU, Indian Telugu in the UK), East Asia (**CHS**, Han Chinese South; **JPT**, Japanese in Tokyo, Japan; **CDX**, Chinese Dai in Xishuangbanna, China; **KHV**, Kinh in Ho Chi Minh City, Vietnam; **CHB**, Han Chinese in Beijing, China) and the Americas (PUR, Puerto Rican from Puerto Rico; CLM, Colombian in Medellin, Colombia; MXL, Mexican ancestry in Los Angeles, California; PEL, Peruvian in Lima, Peru). Codes in bold are populations sampled from their geographic origin, assumed to be less admixed than those sampled from elsewhere, unbolded. Although in the 1000 genomes dataset singleton variants are probably correctly called, to guard against possible sequencing errors and to be maximally conservative singletons were excluded from all analyses. A list of informative bases for the Altai – chimpanzee (PanTro4) – Hg19 alignment was kindly provided by Andrea Manica and Marcos Llorente. Where required, human – chimpanzee alignments were extracted from the Ensembl-Compara eight primate alignments (http://www.ensembl.org/info/data/ftp/). To avoid alignment ambiguities, bases were only accepted if they lay within blocks of at least 300 aligned bases and were 10 or more bases from the nearest gap, even if this was a single base indel.

### Analyses

All analyses were conducted using custom scripts written in C++, available on request. All statistical analyses were conducted in R 3.3.0 (https://cran.r-project.org/). To be conservative, singleton variants were excluded from all analyses. **ABBA-BABA:** Since the 1000 genomes data are low coverage and include much imputations, population allele frequencies are determined with far greater reliability than individual genotypes. Consequently, ABBA-BABA counts were determined probabilistically based on population allele frequencies assuming Hardy-Weinberg. Thus, if allele ‘A’ had frequencies of 0.2 and 0.4 in populations one and two, this site would contribute 0.2 * (1 – 0.4) = 0.12 ABBAs and (1 – 0.2) * 0.4 = 0.32 BABAs, the numbers expected if alleles were drawn at random from each population. **Heterozygosity:** heterozygosities were calculated similarly: if ‘A’ and ‘B’ were at frequencies 0.3 and 0.7, 0.3 * 0.7 * 2 = 0.42 of a heterozygous site was counted, the probability of drawing a heterozygous genotype from that locus if the population were in Hardy-Weinberg equilibrium. **Genetic distance:** since the 1000g data are unphased, genetic distances between haplotypes cannot be calculated. Instead I used an approximation. All variable sites in each individual were recoded as 0, 1 and 2 for ‘AA’, ‘AB’ and ‘BB’ respectively. In pairwise comparisons between individuals, zero and one difference were recorded at sites with the same and different codes, equivalent to assuming that bases that could be identical are identical and that, wherever a difference must exist this always occurs in the most dissimilar haplotype pair. Genetic distance was then taken as the sum of differences within a given window.

### Detecting unusual fragments

For the general analysis, for a given genomic window of 20Kb, all pairwise intra-population distances (PIPD) were calculated (for distance measure, see above). Across the genome, mean PIPD was 45.7 +/-15 s.e.m. To reduce noise, uninformative windows with mean PIPD < 15 were excluded. Each population then yields a maximum divergence and I compared the maximum maximum among the seven African populations with the maximum maximum found across Europe, East Asia and Central Southern Asia. American populations were excluded due the high levels of admixture, though inclusion would have had negligible impact.

To estimate the frequency of objectively unusual fragments in individuals I used a more conservative approach. For the African maximum I used the ‘minimum maximum’, i.e. the smallest of the four maxima across LWK, ESN, MSL & YRI. This guards against the possibility of an occasional Neanderthal fragment in Africa. ASW and ACB were excluded because these populations include appreciable non-African ancestry, while GWD was chosen at random to be reserved as an independent source of control individuals. Each non-African population had one random GWD individual added as an internal control so that allowance could be made for windows with too little differentiation for a Neanderthal sequence to stand out: if the African sequence qualifies as unusual, so too should a more divergent Neanderthal sequence. I further assumed that introgressed fragments were not at high frequency such that any introgressed fragment would yield many (at least 10) comparisons with individuals carrying modern human DNA.

## Acknowledgements

I have had a number of useful and informative discussions Anders Eriksson, Andrea Manica, Mathias Meyer, Rob Foley, Marta Lahr, Eske Willerslev, Martin Sikora, Simon Martin and Mathias Currat. This work was not funded.

